# HyperEx: A Tool to Extract Hypervariable Regions from 16S rRNA Sequencing Data

**DOI:** 10.1101/2021.09.03.455391

**Authors:** Anicet Ebou, Dominique Koua, Adolphe Zeze

## Abstract

The 16S ribosomal RNA gene is one of the most studied genes in biology. This 16S ribosomal RNA importance is due to its wide application in phylogenetics and taxonomic elucidation of bacteria and archaea. Indeed, 16S ribosomal RNA is present in almost all bacteria and archaea and has, among many other useful characteristics, a low mutation rate. The 16S ribosomal RNA is composed of nine hypervariable regions which are commonly targeted by high throughput sequencing technologies in identification or community studies like metabarcoding studies. Unfortunately, the hypervariable regions do not have the same taxonomic resolution among all bacteria taxa. This requires a preliminary *in silico* analysis to determine the best hypervariable regions to target in a particular study. Nevertheless, to the best of our knowledge, no automated primer-based open-source tool exists to extract hypervariable regions from complete or near-complete 16S rRNA sequencing data. Here we present HyperEx which efficiently extracts the hypervariable region of interest based on embedded primers or user-given primers. HyperEx implements the Myers algorithm for the exact pairwise sequence alignment. HyperEx is freely available under the MIT license as an operating system independent Rust command-line tool at https://github.com/Ebedthan/hyperex and https://crates.io.

## INTRODUCTION

For decades, the determination of microbial diversity and their classification has been a major topic of interest (1). To this end, the 16S ribosomal RNA (16S rRNA) gene became a gene widely used in phylogenetics and taxonomy of bacteria (2). It acquired this status because of its ubiquitous presence in bacteria and archaea, its relatively short length (around 1500 bp), its low mutation rate over time, and the fact that it is composed of multiple conserved and variable regions that can be easily targeted thanks to surrounding primers (2, 3). Thanks to ever-improving sequencing technologies, the 16S rRNA has enabled the discovery of novel bacteria and made it possible to study communities of microorganisms in a high throughput manner. However, a large part of the next-generation sequencing tools is only able to sequence short amplicons from 150 bp to 300 bp. Therefore, they are used to sequence portions of the 16S rRNA gene around conserved regions known as the hypervariable regions (4, 5). The 16S rRNA contains 9 hypervariable regions (V-regions) namely V1 to V9 which are bound by a set of primers (4, 6, 7). Depending on the study, the choice of the V-region to be sequenced is of high importance (8). Indeed it has been demonstrated that V-regions taxonomic resolution is not the same among microorganisms (9–12). The taxonomic resolution of the hypervariable region is the ability of a given region to correctly discriminate between microorganisms from different taxonomic groups. Therefore the choice of the V-region is of great interest and remains an ongoing debate (13–15): individual V-regions or their combinations are not necessarily unique for a given taxonomic group. Therefore, selecting a specific V-region is not necessarily sufficient to elucidate internal diversity in a metagenomic-like study. Moreover, a wrong selection of the V-regions can considerably be misleading when assessing the real microbial diversity. For example, we recently demonstrated that the V5-V7 region seems to be more suitable for differentiating rhizobia at the genus level, possibly replacing the common use of the V4-V5 region (8). In consequence, either in metabarcoding or specific bacteria studies using next-generation sequencing tools, an *in silico* analysis for the choice of the best V-region is a necessity. A common preliminary analysis consists in extracting a specific V-region and estimating its discriminatory power toward the taxonomic group of interest by realizing multiple sequence alignments. V-region extraction is done either manually or using V-Xtractor (16). On the one hand, manual extraction is a long process that requires expert knowledge. Even for experts, manual extraction remains a challenge when considering mismatch in primers. On the other hand, V-Xtractor, proposed in 2010, lacks updates of the built-in hidden Markov models and yields sequences requiring time-consuming manual processing. Here we present “HyperEx” to efficiently extract V-regions from sequencing data based on primers sequences. HyperEx is shipped with built-in support for known 16S rRNA primers but can also be used with custom primers. HyperEx stands for HyperVariable Region Extractor.

## MATERIAL AND METHODS

### HyperEx Algorithm

The Myers algorithm is a fast bit-vector algorithm for approximate string matching (17). It uses dynamic programming to rapidly match strings and has an application for pairwise sequence alignment given a distance. HyperEx uses a slightly modified version of the Myers algorithm as implemented by Šošić and Šikić (18), in which base substitution is preferred over base deletion or gap insertion. This implementation, available in the rust-bio package (19), is used by HyperEx to match primers with a given distance (1 by default) which allows mismatches during the matching step. HyperEx takes as input a FASTA/Pearson file either compressed or not and outputs i) the retrieved region in a FASTA file and ii) a General Features File. Submitted nucleotide sequences are searched for every primer in forward and reverse sense. Once found the region between the primers including the primers is extracted and reported.

### Built-in primer selection

We identified the most common 16S rRNA primers as cited in the literature (7, 20). We further numbered the primers based on the *Escherichia coli* 16S rRNA gene system of nomenclature (4, 6) These primers are proposed as part of the HyperEx tool (Table 1).

**Table 1.**
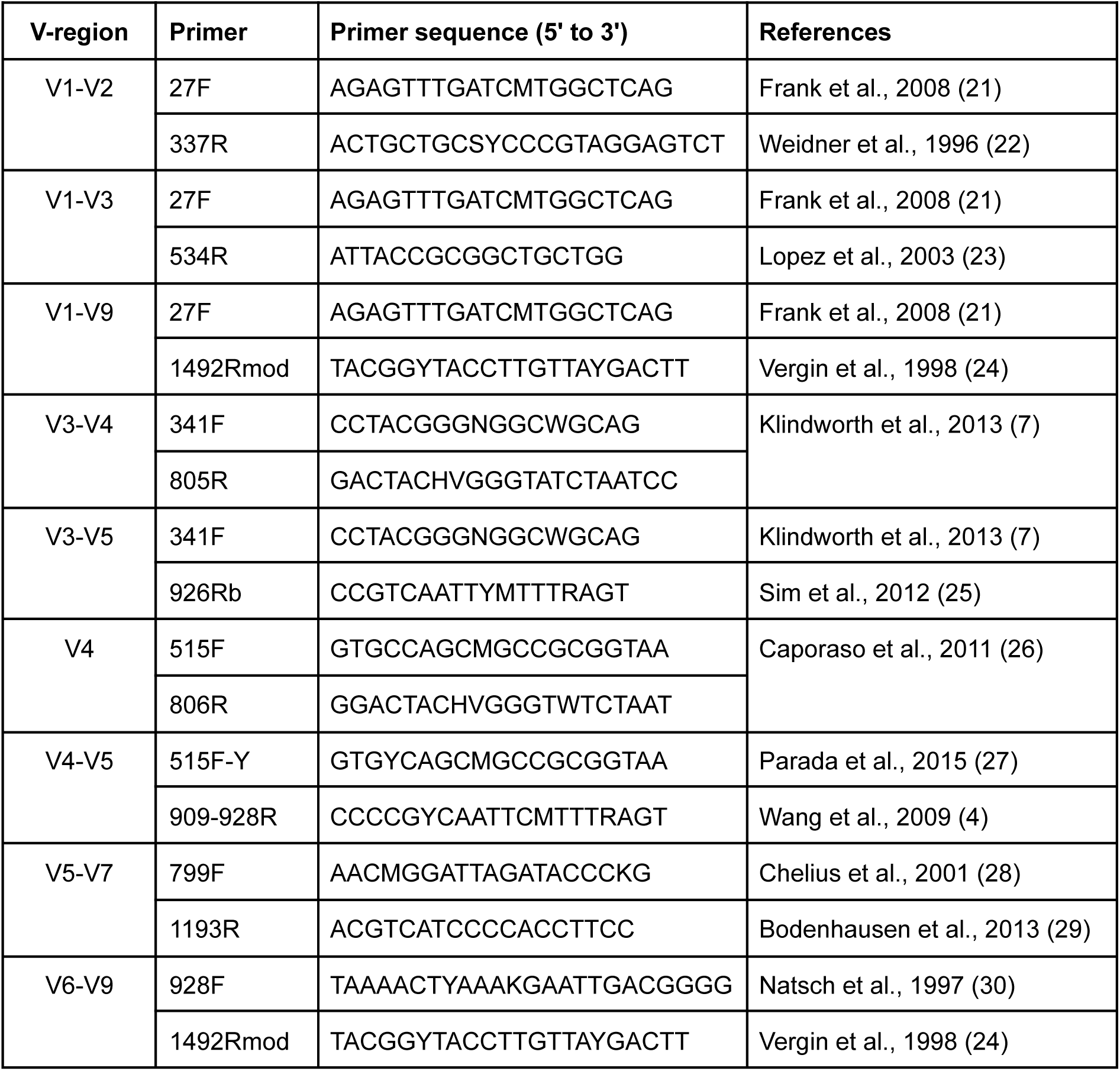
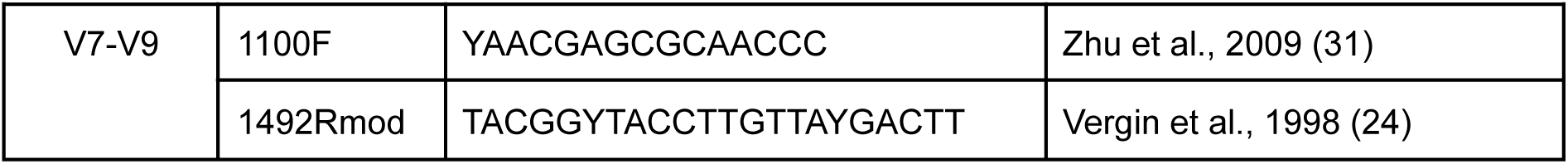
HyperEx built-in primers sequence for 16S rRNA hypervariable regions

### Validation protocol

We performed HyperEx tests on a Linux computer with a 2.70 GHz Intel Core i7 processor and 16 GB RAM and compared HyperEx v0.1 to V-Xtractor v2.1, an HMM-based tool for V-region extraction. The test set was composed of the manually extracted V-region from all ninety-one 16S rRNA sequences of *Rhizobium spp*. as listed on the LPSN (32) (Supplementary file 1). Used sequences have at least 1200 bp and a validly and correctly published name. V-regions were first manually extracted from the sequences using the set of primers defined in Table 1 by searching them in the sequences using Jalvew v2.11.5 (33). Furthermore, to assess the accuracy of tested tools automatically returned V-regions were computationally compared to the manually extracted regions. We conducted an *in silico* investigation of the size of the V-region among *Rhizobium spp*. species using the test set. The 3’- and 5’-end conical structure of *Escherichia coli* 16S rRNA gene described elsewhere (36, 37), together with the annotation and numbering system of *Escherichia coli* K-12 (genome accession number U00096) were used to delineate the full-length size of the 16S rRNA gene sequence.

For accuracy evaluation, we considered only the V3-V4, V3-V5, V4, V4-V5, V5-V7 regions as the used *Rhizobium spp*. 16S rRNA gene data was not complete and misses V-regions V1, V2, and V9.

## RESULTS AND DISCUSSION

### Outline of the proposed HyperEx Tool

The HyperEx tool is written in Rust, was tested on Linux, macOS, and Windows systems, and has precompiled binaries for Linux, macOS, and Windows systems under different processor architecture. It is distributed as an open-source package and full documentation is available on the HyperEx homepage (https://github.com/Ebedthan/hyperex). Once downloaded and installed, one can simply type the following command to extract all V-regions supplying a set of complete 16S rRNA sequences: hyperex <input.fa/input.fa.gz> (values between < and > must be given by the user, for filenames, the file type is indicated even if files could be named without the proposed extension; / stands for “or”).

HyperEx will then search for the supplied or built-in forward and reverse primers for each sequence in the given sequence file using the powerful and fast Myers pattern matching algorithm (Figure 1).

**Figure 1.**
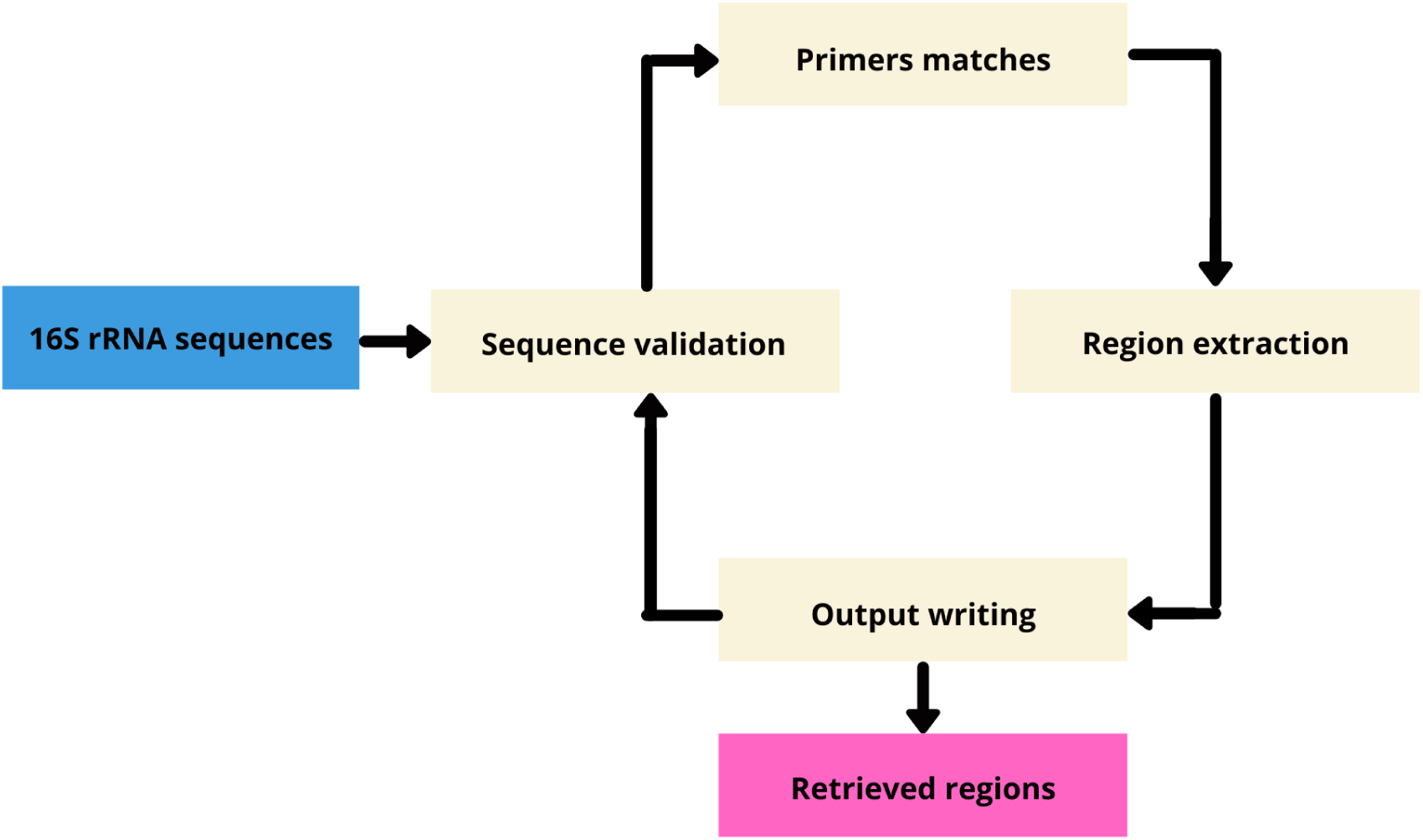
HyperEx processing of 16S rRNA sequences. Input data: Blue, HyperEx processing: Pale brown, Result: Pink.

Supplied primers can either be common 16S rRNA primer sequences (identified by their name or sequences), or custom primers sequences, both with possible IUPAC ambiguities. When a primer file is supplied, built-in ones are not used anymore.

HyperEx allows base pairs mismatch. The user has to indicate the number of accepted mismatches with the mismatch option.

Matching regions (supposedly corresponding to V-regions) are exported in a Fasta/Pearson file along with a Generic Feature Format Version 3 (GFF3) file containing the position of the region in the parent sequence following the specification of the Sequence Ontology (https://github.com/The-Sequence-Ontology/Specifications/blob/master/gff3.md, Accessed on Friday 3 September). Both output files are stored according to the prefix specified or by default in the current working directory.

A report is also provided to allow users to follow the extraction process. Thus, for example, when using built-in primers, when jumping from one to another, if no result was found for a given one, a notice is provided. This can be particularly useful if one is interested in a specific region.

The reproducibility of the region extraction and its speed make HyperEx high throughput compatible and handy.

### Assessing the accuracy and run time of HyperEx

To assess our proposed tool, we compared HyperEx v0.1 to V-Xtractor v2.1, an HMM-based tool for V-region extraction using the test set previously described. Overall, HyperEx consistently extracted the full V-region with 100% of sequence identity compared to predicted V-Xtractor extracted V-regions (Table 2).

**Table 2.** Comparison between the size and sequence identity of the extracted V-regions

| V-region | Expected size | HyperEx extracted V-region |  | V-Xtractor extracted V-region |  |
| --- | --- | --- | --- | --- | --- |
|  |  | size | Sequence identity | size | Sequence identity |
| V3-V4 | 439-442 | 439-442 | 100 % | 232-234 | 52.72% - 53.18% |
| V3-V5 | 560-565 | 560-565 | 100 % | 421-427 | 75.17% - 76.25% |
| V4 | 292-293 | 292-293 | 100 % | 85-86 | 29.10% - 29.45% |
| V4-V5 | 413-419 | 413-419 | 100 % | 274-280 | 66.34% - 67.79% |
| V5-V7 | 412-421 | 412-421 | 100 % | 351-360 | 84.98% - 87.16% |

Moreover, HyperEx completed this task in 0.154 seconds which is more than 58 times faster than V-Xtractor run time.

The difference of accuracy between HyperEx and V-Xtractor can be explained by the fact that V-Xtractor uses HMM models which have not been updated for a while. Moreover, by principle, even when a primer is missing, the HMMs tend to predict distant and incorrect matches. The fast run time shown by HyperEx compared to V-Xtractor is due to the programming language used and the implementation strategy. Although V-Xtractor is fast, its Perl implementation is slower than HyperEx Rust implementation which is a fast, safe and reliable programming language.

HyperEx is, therefore, a reliable tool that can be used in conjunction with tools like RNAmmer (34), rRNASelector (35), or Barrnap (https://github.com/tseemann/barrnap) for automatic *in silico* retrieval of 16S rRNA sequence and 16S rRNA V-regions. Furthermore, HyperEx can also be used with 18S rRNA, 23S rRNA for extraction of their V-regions. To this end, one would just have to provide a file with the appropriate pair of primer sequences.

## SUPPLEMENTARY DATA

Supplementary Data is available at bit.ly/hyperex_sup_file.

## ACKNOWLEDGEMENTS

The authors would like to thank Audrey Y. Addablah and Fossou K. Romain for their valuable comments on the tool and manuscript.

